# Spiking network model of A1 learns temporal filters with frequency preferences

**DOI:** 10.1101/2023.07.10.548413

**Authors:** Danielle Roedel, Braden A. W. Brinkman

**Author notes:** Corresponding author, July 10, 2023.

## Abstract

The sparse coding hypothesis has successfully predicted neural response properties of several sensory brain areas. For example, sparse basis representations of natural images match edge-detecting receptive fields observed in simple cells of primary visual cortex (V1), and sparse representations of natural sounds mimic auditory nerve waveforms. SAILnet, a leaky integrate-and-fire network model (“Sparse and Independently Local network”) has previously been shown to learn simple V1 receptive fields when trained on natural images. Experimental work rewiring visual input to auditory cortex found that auditory neurons developed visual response properties, suggesting that developmental rules may be shared across sensory cortices.

In this work we adapt SAILnet to train it on waveforms of auditory sounds and learn temporal receptive fields (filters), in contrast with previous work that trained SAILnet or other network models on spectrograms. In our model network of primary auditory cortex (A1) neurons receive synaptic current from input neurons who temporally filter the direct sound waveforms. To show the network learns frequency-dependent filters naturally, we do not parametrize the temporal filters, and only restrict the total number of time points in the filters. To make training feasible, we simplify the model to a single input neuron and 768 A1 neurons, and we train the network on “lo-fi” music, whose spectral power is limited to frequencies of *∼*10, 000 Hz or less, giving a manageable temporal resolution of the stimulus and filters. The learned filters develop distinct frequency preferences, and reconstruction of novel stimuli captures the low-frequency content of signals in reasonable detail, with audio playback capturing clear aspects of the original stimulus. Lastly, our work also has a pedagogical benefit: the learned stimulus features can be played as sounds, which aids in teaching sensory coding to learners with visual impairments who cannot perceive stimulus features learned by V1 models.

## Introduction

A foundational goal of theoretical neuroscience is understanding the rules by which sensory cortices develop and represent different types of stimuli. The sparse coding hypothesis provides an information theoretic explanation, with metabolic interpretations, for these developmental rules. This hypothesis suggests that only a small number of neurons will be firing at once (population sparseness), or that each neuron is only responsive to specific stimuli so that it does not fire very often (lifetime sparseness) [1]. This hypothesis has motivated network models of primary visual cortex (V1) that learn simple cell receptive fields similar to those those found experimentally [2–4].

Following foundational work by Olshausen and Field [3], Zylberberg et al. [1] proposed “SAILnet” (for “Sparse and Independent Local network”), a spiking network model with biologically plausible learning rules model that developed sensitivity to visual features statistically similar to receptive fields experimentally observed in simple cells in visual cortex. Similar to the model of [3], the learning rules in SAILnet are motivated by sparse coding theory. One learning rule, for example, promotes sparseness in firing to mimic homeostatic mechanisms in the brain. Another minimizes correlations in neural activity, which is similar to the effects of lateral inhibition present in cortical neurons. Furthermore, these learning rules function locally, meaning they change parameters based only on activity of the pre- and post-synaptic neurons from a single synapse, rather than the activity of the entire network.

While SAILnet and related theoretical work have been developed to understand and explain sparse coding in the visual system, experimental work has suggested that similar learning rules may also apply in the primary auditory cortex (A1). In particular, Roe et al. [5] rewired ferret retinal projections into the auditory thalamus, bypassing the visual thalamus. Following this rewiring and presentation of visual stimuli, A1 cells exhibited orientation and direction selectivity, similar to simple cells found in V1 [5]. Furthermore, experimental work has also shown that A1 neurons in unanesthetized rats exhibit population sparseness in response to various auditory stimuli [6].

Training network models on auditory waveforms is more difficult than visual stimuli, whose temporal properties can often be neglected (e.g., during periods of fixation). Many studies have circumvented this difficulty by training networks on visual representations of sounds, including spectrograms and cochleograms [7–11]. These representations mimic the pre-processing of auditory stimuli by the early auditory pathway, and most A1 models are trained on this pre-processed input to study the emergence of “spectrotemporal” stimulus feature preferences, similar to those observed in the inferior colliculus, auditory thalamus, and A1 [12]. Models trained on raw auditory sound waveforms are usually intended as models of auditory nerve fibers in the early auditory pathway [13], rather than any firing observed in downstream structures.

In this work, our goal is to modify the SAILnet model of spiking neurons and train it on raw auditory waveforms. Because the recurrently connected neurons in the network comprise a model of primary auditory cortex, the temporal receptive fields learned by our network serve as a simplified model of the early auditory pathway, bridging the two types of former approaches that modelled either early auditory nerve fiber responses or A1 responses, but not both. We show that our model neurons learns temporal receptive fields with distinct frequency preferences without using basis functions, and reconstruction of novel stimuli captures the low frequency content of signals in reasonable detail. This demonstrates that our model learns information-rich features of signals, providing theoretical evidence that sparse coding rules used in models of V1 can also apply to models of A1.

## Results

### Modified SAILnet as a model of A1

Our network model is a modified version of SAILnet (Sparse and Independent Local network), which comprises a spiking, leaky integrate-and-fire network model of primary visual cortex (V1) [1]. To study neural coding of auditory stimuli in A1 we modify SAILnet to receive auditory input in the form of sound waveforms *X_t_*, as shown in Fig. 1. The network consists of *N* neurons whose membrane potentials evolve according to the discrete-time dynamics

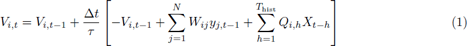

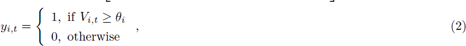

followed by a reset

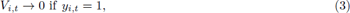

**Figure 1:**
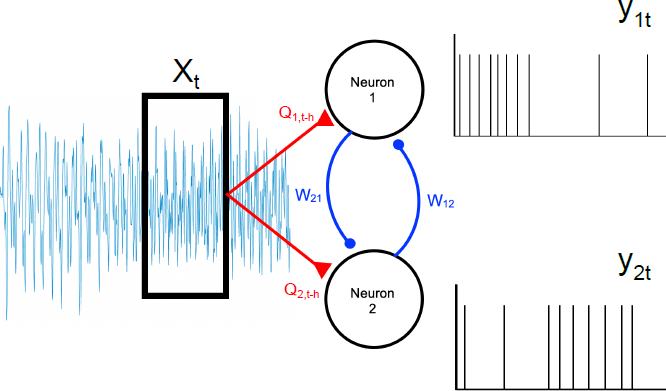
Schematic of the network model showing two model neurons. Our primary auditory cortex (A1) network is a modified version of SAILnet, comprising a network of leaky integrate-and-fire neurons [1]. Our 768 neurons (two representative neurons shown above) receive auditory input in the form of randomly-drawn clips from lo-fi music sound waves (*X_t_*, selected clip from longer audio) which are filtered by each respective neuron’s receptive field, *Q_i,t−h_* with a time lag *h*. Each neuron receives inhibitory input from all other neurons with strength *W_ij_*. Each neuron *i* generates a spiking output at each time point *t*, *y_it_* = 1 if the neuron spikes and 0 otherwise. The input weights *Q*, lateral weights *W*, and firing threshold of each neuron *θ_i_* (not depicted) are learned by training the network on lo-fi audio clips.

where *V_i,t_*is the membrane potential of neuron *i* at time-step *t*, Δ*t/τ* = 0.1 is the size of the time-step relative to the membrane time-constant, *W_ij_* is the strength of the synaptic connection from neuron *j* to neuron *i* (constrained to be negative, modeling the effective mutual inhibition of excitatory principal neurons through inhibitory interneurons), *y_j,t−_*_1_ is the number of spikes (0 or 1) fired by neuron *j* at the previous time-step, *θ_i_* is the firing threshold of neuron *i*, and *Q_i,h_* is the stimulus filter from the input neuron to neuron *i* at time lag *h*, which weights the past history of the stimulus *X_t−h_*up to *h* = *T*_hist_ = 400 time-steps before the current time-step *t*.

Most investigations of neural coding in A1 assume that the early auditory pathways pre-process sound waveforms, converting them into “spectrograms,” which plot the power of the signal as a function of frequency at different moments in time throughout the auditory signal. Experimentally observed A1 neuron responses can be modeled as a convolution between a “spectro-temporal receptive field” and the spectrogram of the auditory input [7–12]. In contrast, in this work we train our A1 network on the sound waveforms directly. In doing so we are effectively modeling part of the development of the spectrotemporal receptive fields: if we consider a collection of *N*_input_ input neurons, then the stimulus filters *Q_ik,h_*—the filter from input neuron *k* to cortical neuron *i* at time-lag *h*—can be interpreted as spectrotemporal receptive fields when the input neurons are sorted according to the dominant frequency their corresponding input filter. In this study we only consider a single input neuron to render the training computationally feasible, but in future studies the addition of multiple input neurons will allow our approach to make contact to spectral models of auditory coding.

The sound waveforms we use for training comprise royalty-free lo-fi music. This choice of auditory stimulus is motivated by selecting stimulus statistics with relatively low frequency content, as natural sounds tend to have very high frequencies, which would require very fine temporal resolution— small time-steps in the model—which is computationally intractable unless some method of dimensionality reduction is performed on the training stimuli. The only stimulus pre-processing we perform is a downsampling of the songs from 44,100 Hz to 2,000 Hz and a normalization of the songs to have zero mean and unit variance, similar to the preprocessing done on the training images for the visual SAILnet [1, 9]. The original lo-fi songs vary in length, so we take 60 seconds of each song to use as a training set. For each training iteration of the network, 10 clips of 384 milliseconds (768 time points at a sampling frequency of 2, 000 Hz) are randomly drawn from this training set to feed to 10 copies of the network in order to train the network. The input currents generated by these auditory stimuli appearing in Eq. (1) correspond to the term *^T^*^hist^ *Q_i,h_X_t−h_*. The input filters *Q_i,h_*, recurrent synaptic connections *W_ij_*, and the firing thresholds *θ_i_*are trained in response to presentations of these auditory stimuli. The dynamics of these parameters are Hebbian-like synaptic and homeostatic learning rules inspired by sparse coding theory. The learning rules promote minimal correlation in spiking (Eq. (4)), sparseness in firing (Eq. (5)), and linear decoding of the auditory signal from the emitted spike trains (Eq. (6)):

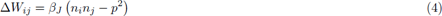

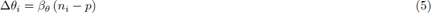

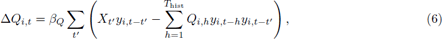

where Δ*Q_i,t_*, Δ*W_ij_*, and Δ*θ_i_* are the updates to the network parameters after a series of stimulus presentations, the *β*’s are learning rates, 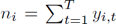 is the total number of spikes fired by neuron *i* over the course of *T* time-steps of the auditory input, and *p <* 1 is the target number of spikes each neuron should fire in this window. The first two learning rules are unaltered from the visual SAILnet model, while the third learning rule is modified to account for our auditory stimuli. Specifically, the network is designed to approximately linearly encode the auditory stimuli in the spiking activity of the neurons: *X_t_* ≈ Σ*_i,h_ y_i,t−h_Q_i,h_*. The learning rule is then derived by gradient descent of a mean-squared error between the stimulus and this linear decoding, neglecting non-local terms, as in the visual SAILnet model [1].

We train our network for 9, 000 total iterations to test convergence and overfitting. The optimal level of training appears to occur around 6000 *∼* 7500 iterations (see Quantification of Model Performance), so for the rest of this work we focus on the network obtained by 6, 000 iterations, unless otherwise specified.

### SAILnet learns temporal receptive fields with diverse frequency preferences

The 768 neurons in our network learn temporal filters with distinct frequency preferences (Fig. 2). In contrast to other studies that have constrained their training using basis functions [8, 9], the only constraint used in our training was the number of time points in each temporal filter, so the oscillatory temporal filters are naturally learned by the model and not built in.

**Figure 2:**
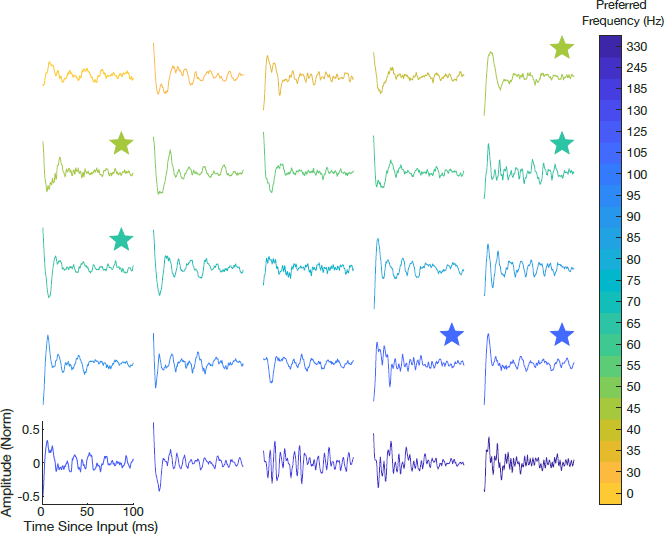
Temporal receptive fields learned by auditory SAILnet. Each of the 768 units comprising our network learns a temporal filter; a subset of 25 temporal filters is shown above. Each box represents one unit’s temporal receptive field. The only constraint on the learning of these filters is the temporal resolution (number of time points). We use a Fourier analysis to determine the preferred frequency of each temporal filter, binning the results into 22 unique preferred frequencies (Fig. 4). The figure above shows two randomly selected receptive fields from the 3 most common unique preferred frequencies (45, 65, and 105 Hz) and one randomly selected receptive field from all other unique preferred frequencies.

While lo-fi music typically consists of frequencies up to 10, 000 Hz, spectral analysis reveals that our training audios have no significant frequency components over 1, 000 Hz (Supplementary Fig. 10). For this reason, we take 1, 000 Hz to be the Nyquist frequency and downsample all ten songs to 2, 000 Hz. We therefore expect the frequencies that our model can learn to be restricted to this range. We find the majority of the frequency components from all ten downsampled and normalized training audios were 400 Hz or less (Fig. 3).

**Figure 3:**
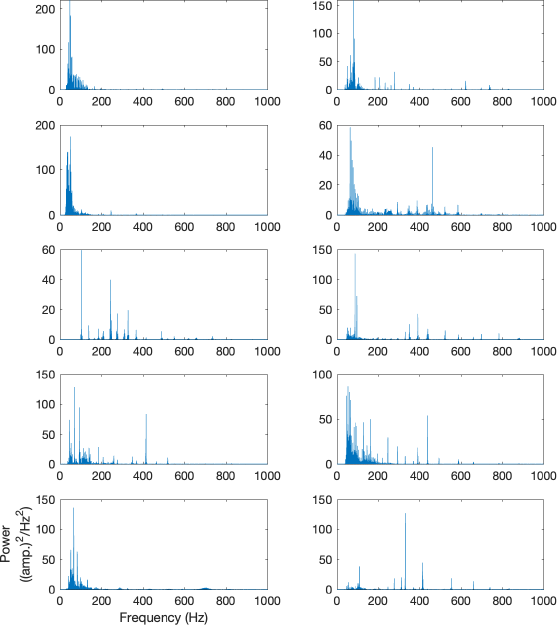
Power spectra of downsampled and normalized training sounds. We downsample the lo-fi audio clips used during training (Supplemental Fig. 10) to 2,000 Hz and normalize them to have zero mean and unit variance. We take a 60 second clip from each of these processed training waveforms, of which the power spectra are shown above. Most of the frequency components are 400 Hz or less, so we expect the network to capture these frequency components in the most detail.

To assess the preferred frequency of each neuron we compute the discrete Fourier transform for each of the 768 learned temporal filters and identify the frequency with the greatest spectral power. The preferred frequencies ranged from 0 Hz to 330 Hz (Fig. 4). This does not cover the full range of frequency components observed in our training audios, but does match the most frequently observed frequency components (Fig. 3).

**Figure 4:**
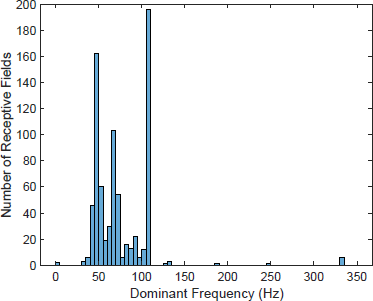
Histogram of preferred frequencies of learned temporal filters. We compute the discrete Fourier transform for each of the 768 neurons in our network and define a neuron’s preferred frequency as the frequency with the greatest power. After binning there are 22 unique preferred frequencies displayed in the histogram above. These preferred frequencies vary from 0 to 330 Hz with 98% of them lying between 30 Hz and 110 Hz.

### Reconstructions of novel stimuli capture low-frequency content of signals

We investigate how well our trained networks represent auditory stimuli by simulating network activity in response to ten novel lo-fi song inputs that were not part of the training set and decoding the spiking activity to predict the stimulus. For the sake of conciseness, in this section we focus on the reconstruction of one novel audio (Fig. 5). We present the normalized and downsampled audio to the network in patches of 768 time-steps (384 ms) and record the network’s response to the stimuli. We then convolve this network activity with the learned stimulus filters to decode the stimulus. Because SAILnet only predicts deviations from a baseline, we add the mean and variance of the original stimulus back to the decoded signal to reconstruct the novel lo-fi audio. Low frequency content up to approximately 200 Hz was captured in reasonable detail (Fig. 6). With audio playback, aspects of the original sound are clearly present in the reconstruction. These frequencies captured in the reconstruction coincide with the preferred frequencies of the temporal filters (Fig. 4). The higher frequency content of the signals is not captured well by our trained network because few neurons developed stimulus filters sensitive to such high frequencies.

**Figure 5:**
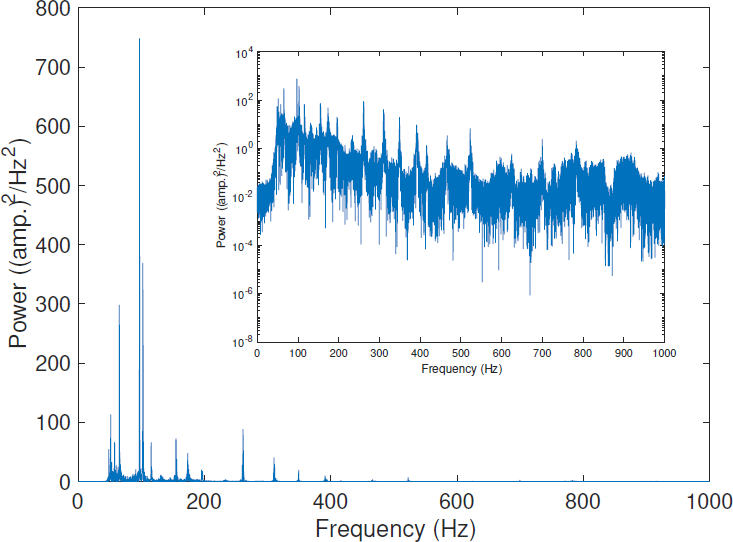
Power spectrum of a representative novel lo-fi sound clip. obtained from pixabay.com, plotted on linear and log scales (inset). Similar to the lo-fi songs used for training, there are no significant frequency components above 1,000 Hz. Following downsampling and normalization, the majority of the frequency components are less than 400 Hz. The frequency components of this novel audio are similar to the frequencies comprising the training clips, shown in Fig. 3.

**Figure 6:**
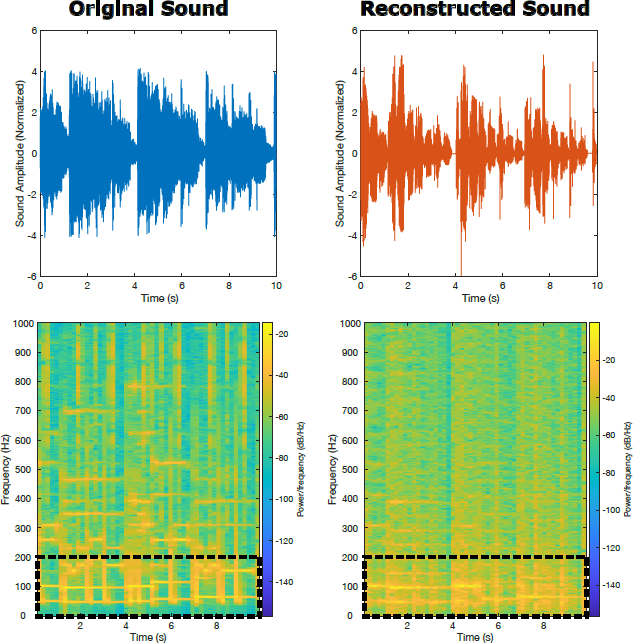
Visual representations of a novel lo-fi audio and the reconstruction of that audio. **Left:** Waveform (top) and spectrogram (bottom) of a novel lo-fi audio clip downsampled to 2,000 Hz. The normalized clip is input to the trained network to generate spiking activity, which is then decoded and reconstructed. **Right:** Waveform (top) and spectrogram (bottom) of the reconstructed audio. High frequency attenuation present in the original waveform is captured in the reconstructed waveform. The low frequency content (up to *∼*200 Hz) of the original signal is captured in reasonable detail, which is illustrated in the spectrograms. Higher frequency features of the sound are not captured by our network because few neurons developed stimulus filters sensitive to such high frequencies (Fig. 4).

### Quantification of model performance

To quantify the trained network’s performance we calculate several measures of similarity between the true and reconstructed stimuli at seven different points throughout training, from iteration zero (before any training) to iteration 9,000, every 1,500 iterations (representative examples shown in Supplementary Figs. 11, 12). For each reconstruction, we calculate i) the cross-correlation of the original signal and its reconstruction to find the lag at which the correlation between the two signals was maximized, ii) the point-wise mean squared difference between the temporal waveforms of original signal and the shifted reconstructed signal (i.e., the reconstructed signal shifted by the lag found in i), and iii) the point-wise mean squared difference between the spectrograms of the original signal and the shifted reconstructed signal; see Methods and Supplementary Figures showing representative examples of how the waveforms and spectrograms evolve during training, Figs. 11, 12).

We find the peak correlation increases rapidly between iterations 0 and 1500, plateauing at a value of 0.4 for the remaining iterations (Fig. 7A), indicating a reasonable similarity between the original and reconstructed signals. As can be seen in Fig. 6, the reconstructed signal shows some attenuation due to reconstructing the signal in several patches, contributing to some of the dissimilarity between the two signals. Next, we calculate the point-wise mean squared difference between the original signal and the reconstructed signal, shifted so that we are comparing the two signals when they are maximally correlated. If we do not control for this phase offset the point-wise mean squared error is not a reliable measure of the similarity because of destructive interference between the signals (Supplementary Fig. 9). While the mean squared error decreases slightly with training, the decrease appears modest by this measure, dropping to only 85% of the mean squared error of the untrained network (Fig. 7B).

**Figure 7:**
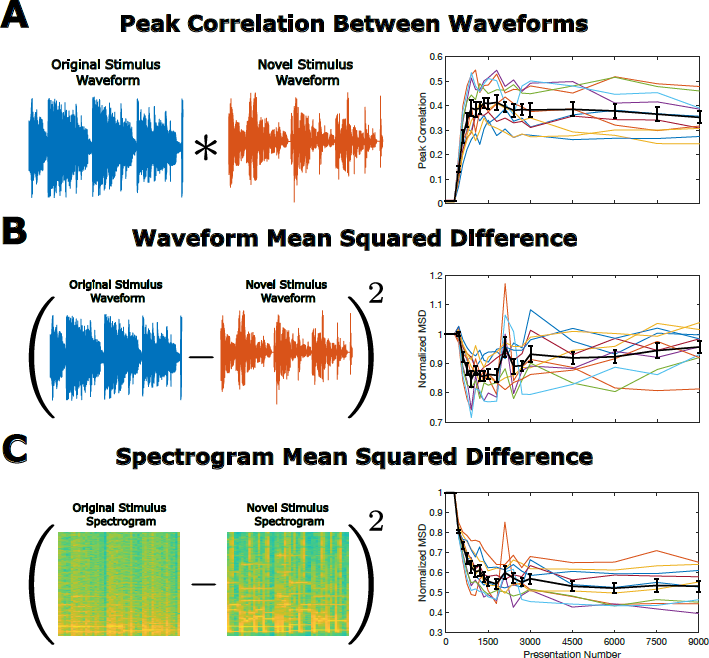
Quantification of model performance. A). Peak correlation between waveforms. For each reconstruction of a novel lo-fi audio clip we calculate the cross-correlation to determine the time lag at which the original signal and the reconstructed signal are most similar. The above plot shows the maximum correlation between the original signal and the reconstruction for 10 novel lo-fi audio clips (colored lines) and the mean *±* standard error of the set (black line). There is an initial increase from iteration 0 to iteration 1, 000 followed by a plateau at a correlation value around 0.4. **B. Waveform mean squared difference:** Point-wise mean squared difference between the waveform of the novel lo-fi audio and reconstructed waveform shifted by the time-lag to compare the maximally correlated signals. This metric exhibits a slight decrease, indicating some improvement over the untrained network. **C. Spectrogram mean squared difference:** For each reconstruction of each novel lo-fi audio, we calculate the mean squared difference between the spectrograms of the original sound and the reconstructed sound. The above plot shows this metric 10 novel lo-fi audios (colored lines) and the mean *±* standard error of the set (black line). This metric sharply decreases between iterations 0 and 3,000, and then plateaus.

Finally, we calculate the mean squared difference between the spectrograms of the original and recon-structed signals. This measure decreases to about 55% of the mean squared difference between the original spectrogram and the reconstruction using the untrained network (Fig. 7C), despite the fact that only the low frequency content of the spectrogram appears to be captured well by our trained networks.

All together, the patterns observed in these three metrics of network performance show that training results in reconstructions that capture many features of original signal, quantifying the qualitative similarity that can be heard by comparing playback of the original and reconstructed sounds (Supplementary Files 6a and 6b). This demonstrates that our network successfully learns information-rich features of auditory signals.

## Discussion

Our results demonstrate that the Sparse and Independent Local network (SAILnet) model, originally developed to learn simple cell receptive fields observed in primary visual cortex, can be modified to learn temporal receptive fields when trained on auditory sound waves. We train our model on lo-fi music, which only contains frequency components up to 10,000 Hz. In response to this training, our 768 model neurons learn temporal receptive fields with oscillatory components at frequencies in the range of 0 to 330 Hz, though 98% of the neurons exhibited preferred frequencies between 30 and 100 Hz (Fig. 4). Notably, we do not impose any constraints on the learned temporal filters other than the number of time points in each filter, so the learned sensitivity to different frequencies is a consequence of the stimulus statistics and learning rules. Following training, we present the network with ten novel lo-fi songs and performed linear decoding of the neuronal activity to reconstruct the stimulus. Aspects of the novel stimuli are captured with audio playback of the reconstruction, and spectrogram analysis shows that low-frequency content of the sound is reconstructed in reasonable detail (Fig. 6). Further quantitative analysis reveals that peak correlation between the waveforms increases with training, and the mean squared differences between both waveforms and the spectrograms decrease with training (Fig. 7), demonstrating that our network successfully encodes auditory information.

### Comparison to prior results and implications for future experiments

As the learning rules used in SAILnet are motivated by information theoretic principles constrained by biologically plausible restrictions on locality, one might hope that similar learning rules should hold in different sensory cortices, with some minor adaptation of the rules to account for differences between the different types of stimuli. Support for this hypothesis is provided by experimental work rewiring ferret retinal projections to the auditory thalamus [5]. With visual input, the neurons in the rewired auditory cortex (A1) exhibited receptive fields similar to those seen in simple and complex cells of the primary visual cortex (V1), indicating the development of these two sensory cortices could be modeled using similar rules. Other experimental work with unanesthetized rats demonstrated that A1 neurons exhibit population sparseness in response to tones, sweeps, white noise bursts, and natural sounds, indicating the sparse coding hypothesis could apply to A1 as well as V1 [6]. These experimental works motivated the present study.

Many prior theoretical studies have also investigated sparse coding in a variety of models. Dodds & DeWeese [9] also modified SAILnet to train on auditory stimuli, but trained on spectrograms of speech, rather than sound waveforms. In their work, [9] employed basis function representation and other methods to reduce the dimensionality of the stimuli. For example, they whitened their stimuli by normalizing the first 200 components resulting from their principal component analysis (PCA) and discarding the remaining components at all. Notably, Dodds & DeWeese found that SAILnet was most successful when stimuli were normalized by removing the mean and variance: differences in how variance is accounted for can alter the population sparseness and the features detected by SAILnet. Such whitening is thought to be performed by the retina and lateral geniculate nucleus in the visual pathway, but it is not clear that a similar procedure occurs in the auditory pathway.

Many studies prior to the development of the SAILnet model have also trained sparse coding models on spectrograms and cochleagrams [7–11]. In [8], for example, spectrograms depicted speech as a function of time and frequency, while the cochleagrams represented speech after it had been processed by the cochlea. This model learned features that were observed in spectrotemporal receptive fields throughout the mammalian visual pathway including the inferior colliculus, auditory thalamus, and A1. This work, however, did not implement any biological constraints and relied on the statistical features of the training sounds to train the model. Ref. [8]’s model learned spectrotemporal receptive fields that also exhibited frequency ranges much lower than those observe in mammals, concluding that this was due to the nature of training only on human speech, which consists of predominantly low frequencies. This accords with our observations that our model’s learned receptive fields developed preferred frequencies between 30 Hz and 110 Hz (Fig. 4), which is very low compared to the range of up to 20,000 Hz that can be heard by humans [14] but coincides with the frequencies of our training audios (Fig. 3).

Some previous studies have used auditory waveforms as training stimuli in models of auditory nerve firing upstream of A1 circuitry [13]. The input neuron in our model can be interpreted as a putative nerve-cell, and thus our auditory SAILnet model can be viewed as a simplified model the encapsulates both the early auditory pathway and A1. Future work will increase the number of input neurons to obtain a better model for the early auditory pathway in our SAILnet adaptation. Such a collection of input neurons can be organized according to their frequency preferences, and in this way would constitute a spectrotemporal receptive field for each neuron.

### Limitations of the model

One of the major challenges for training networks on sensory signals is the high dimensionality of the stimuli in question. In the case of auditory stimuli, this dimensionality comes in the form of the temporal resolution of the of the sound waveform. The temporal resolution sets the maximum frequency that can be represented, such that representing a 10 kHz sound would require a time-step of 0.05 ms, taking into account the Nyquist frequency. In SAILnet the number of neurons in the network is taken to be greater than the dimensionality of the stimulus in order for the network to learn an “overcomplete basis,” which allows multiple neurons to represent similar stimulus features, bestowing the network with robustness to losing some neurons.

The high dimensionality of stimuli necessitates some form of dimensionality reduction in order to render training tractable, as the computational costs of training extremely large models is prohibitive. As we wanted to determine whether our auditory SAILnet model would learn frequency preferences, we did not want to use basis function representation or PCA-based methods like [9], only the temporal stimulus input. In order to make such training manageable, we opted to downsample our auditory signals so that the highest frequencies represented would be 2, 000 Hz, meaning a time-step size of 0.1 ms requires 768 points to represent a 384 ms clip. We also used a 1 : 1 ratio of neurons to stimulus points (an overcompleteness ratio of 1), the minimum that can be used for an overcomplete basis.

In order to not lose important stimulus features during this downsampling, we had to identify a stimulus set that does not typically contain frequency content above 2, 000 Hz. While natural sounds would be ideal to train the model on, as an auditory analogue of training visual SAILnet on natural images, natural sounds contain significant frequency content up to 20 kHz, too large for our model. We ultimately opted for lo-fi music, obtained from royalty-free music websites, which is typically limited to frequencies of 10, 000 Hz or less. While these lo-fi songs originally had sampling rates of either 44, 100 Hz or 48, 000 Hz, Fourier analysis revealed that there were no significant frequency components greater than 1,000 Hz, allowing us to choose 2, 000 Hz as a Nyquist frequency and downsample to this resolution without sacrificing too much of the original signal.

Related to the computational costs of training, another aspect of our model that we had to limit was the number of input neurons. Our current model only accounts for a single input neuron, and as such only one small clip of a sound provides input to the A1 neurons in our model. Biologically, however, A1 neurons do not simultaneously receive the same auditory signal. An auditory stimulus arriving in A1 could be delayed due to the position of the sound (if it is closer to one ear versus another) or depending on which portion of the cochlea is excited by the sound. Increasing the number of input neurons would allow each input neuron to filter different auditory inputs (which could be the same signal but delayed in time for each input neuron), which could allow for better reconstructions of the auditory signal, and to construct spectrotemporal receptive fields, as noted earlier.

### Pedagogical motivations for this study

In addition to our scientific goal of extending SAILnet to include such a model of the early processing pathway, one of the motivations for encoding sound waveforms directly is pedagogical: the learned receptive fields can be played directly as sounds (Supplementary files 1-5), facilitating teaching of sensory coding concepts to blind or low vision students (BLV). There are 3.8 million BLV adults in the United States working on a post-secondary degree [15,16]; while data on BLV student success in higher education is limited [17,18], some studies have found that BLV students tend to favor majors in humanities subjects over STEM fields [19, 20].

The visual nature of scientific data presentation can be an obstacle for BLV students interested in topics like mathematical model, simulation, and neuroscience. The receptive field is a fundamental concept in neuroscience, and is often introduced through the responses of neurons in the visual cortex. These receptive fields are often represented as two-dimensional heat-maps, whose structure cannot clearly be seen by some BLV students. While in-person classes can sometimes overcome such obstacles by representing plots using tactile 3d printouts [21, 22], this means of non-visual communication is not suitable for digital media, such as online classes or outreach material available online. In some cases sonification of data can be used to communicate information contained in plots [23], but for representing complex data like sensory receptive fields it would be more natural to directly play an auditory receptive field rather than a sonified version of a visual receptive field. In addition to the scientific contribution of this paper, we hope the model and results of this study will be incorporated into computational neuroscience lectures on sensory coding, improving the accessibility of this material for students with limited vision.

## Methods

### Network model

Our model is a modified version of the Sparse and Independent Local network (SAILnet), a leaky integrate-and-fire model of V1 [1]. The original SAILnet and our modified network are written in MATLAB and all subsequent analyses and experiments were performed in MATLAB. SAILnet consists of *N* neurons whose membrane potentials evolve according to the discrete-time dynamics given in the main text (Eqs. 1-3). While the original SAILnet was trained on visual stimuli to model V1, our modified network trains on auditory stimuli to model A1. As shown schematically in Fig. 1, each of the *N* neurons receives auditory input in the form of sound waveforms *X_t_*. Our model structure includes *N* = 768 neurons, which constrasts the 1,024 neurons of the visual SAILnet [1]. Training our model at that scale would have been too demanding in terms of run time and computational power, so we balanced computational costs by setting *N* = 768 as a halfway point between 2^9^ and 2^10^. We set the target average number of spikes fired in response to a stimulus to *p* = 0.05, the value used in the visual SAILnet [1]. Because we train our network for 768 time steps (the length of the auditory stimulus), in contrast to 50 time-steps in the visual SAILnet model, this amounts to a stronger sparsity penalty on the firing of the auditory neurons in our network model.

The network learns stimulus filters *Q_i,h_*, synaptic weights *W_ij_*, and thresholds *θ_i_* according to the Hebbian-like synaptic and homeostatic learning rules stated in the main text (Eqs. 4-6). These rules are inspired by sparse coding theory, and promote minimal correlation in spiking (Eq. 4), sparseness in firing (Eq. 5), and linear decoding of the image patches from the spike rates (Eq. 6). We conducted an early exploratory investigation of the parameter set and used the parameters summarized in Table 1 for our final network, which takes about 28 hours to train on a server of 128 CPU cores (AMD EPYC 7662 2 GHz) with 512 GB of RAM.

**Table 1:** Parameters and standard values used in this work. This table summarizes the parameters that appear in the network equations (Eq. 1 - 3) and the learning rules (Eq. 4-6).

| Parameter | symbol | value |
| --- | --- | --- |
| Time-step relative to membrane time-constant | $\Delta t/\tau$ | 0.1 |
| Number of time points in temporal receptive field $Q_{i,h}$ | $T_{\text{hist}}$ | 400 |
| Target number of spikes fired over stimulus window | $p$ | 0.05 |
| Synaptic weights learning rate | $\beta_J$ | 1 |
| Stimulus filters’ learning rate | $\beta_Q$ | 0.01 |
| Threshold learning rate | $\beta_\theta$ | 0.1 |
| Number of time points in stimulus | $T$ | 768 |
| Number of neurons | $N$ | 768 |

### Training the network

The SAILnet model of V1 was trained on natural images [1], so we initially used natural sounds as analogous stimuli to train our modified model of A1. These natural sounds contained significant power at high frequencies, which was distorted when we downsampled these sounds to a lower sampling rate. Training on these audios without significant downsampling was not computationally feasible. In order to demonstrate that the auditory SAILnet model learned plausible receptive fields, we trained the network on lo-fi music, a genre of music known to have few if any frequency components above 10, 000 Hz. We obtained 10 lo-fi songs from royalty-free music sites including pixabay.com, tunetank.com, chosic.com, and free-stock-music.com. These songs varied in length from 70 seconds to 240 seconds and were all originally sampled at 44,100 Hz. To get a better understanding of the frequency components comprising our training audios, we computed the Fourier transform and plotted the power spectrum as a function of frequency [24]. This analysis confirmed these training audios had no significant frequency components above 1,000 Hz, which we took to be our Nyquist frequency (Supplemental Fig. 10). As in the visual SAILnet model [1], we normalized these downsampled sounds to have zero mean and unit variance, an operation which appears to be necessary for SAILnet’s performance [9]. To account for the varying lengths of the songs, we used 60 seconds of each song as our training set. All song files had two channels, so we took these clips from channel 1 in all cases.

At the beginning of training, all synaptic connection strengths *W_ij_* are set to zero, all firing thresholds *θ_i_* are set to 2, and all stimulus filters, *Q_i,h_* are randomly initialized from the standard normal distribution for each time point *h ∈ {*1, …, 400*}*. We trained the model on sound clips of 768 time points (384 ms) in length, randomly drawn from the bank of 10 longer sound clips. For each iteration of training, batches of 10 randomly selected sound clips are presented to the network in parallel. The network spiking activity obtained in response to all ten sound clips is averaged across the batch and used to update the network parameters according to the learning rules (Eqs. 4-6). The original visual SAILnet trained on stimuli in batches of 100 [1]; our auditory SAILnet model with 10 batches took approximately 28 hours to train for 9,000 iterations on a server of 128 CPU cores (AMD EPYC 7662 2 GHz) with 512 GB of RAM, so a ten-fold increase in batch size was not computationally feasible.

While we trained the network trained for 9,000 iterations, the results we presented in this work are from the network at 6,000 iterations of training. The measures of model performance, including correlation between waveforms and mean squared difference between spectrograms level off by 6,000 iterations (Fig. 7), and while further iterations do not appear to degrade network performance substantially the chances of repeated stimuli overlaps increases with longer training, allowing for the possibility of some over-training by 9, 000 iterations.

### Spectral analysis

We perform two types of spectral analysis in this work. We first computed the discrete Fourier transform of the training audios and plotted the power spectrum as a function of frequency [24]. This was also done for the novel audios. We also computed the discrete Fourier transform of each of the learned temporal filters to find the frequency components of the filters [25]. The most prevalent frequency component was considered the preferred frequency of that stimulus filter.

### Sound reconstructions

Following training we assessed the quality of the network’s representation of the auditory stimulus by simulating network activity in response to ten novel inputs and decoding the spiking activity to predict the stimuli. Our novel inputs were lo-fi songs obtained from the same royalty-free websites as our training audios. These songs were not a part of our training set, but did have similar frequency components to the songs in our training set (Figs. 5, 10). As we did with our training audios, we downsampled these songs to 2,000 Hz and normalized them to have zero mean and unit variance and used only the first of two audio channels.

We shortened each novel lo-fi song to contain 19, 968 time points (9.984 seconds) so that the clips could be presented to the network in 26 non-overlapping patches of 768 time points (384 ms). This patch length was chosen to match the length of the audio clips used for training the network. We normalized each patch to have zero mean and unit variance and presented them to the network one at a time, recording the network activity in response to each patch. Presenting the novel audio to the network using this patchwise method coincides with presenting the novel image to the visual SAILnet in patches of pixels that are normalized to have zero mean and unit variance [1].

To decode the original sound, we convolved the neuronal activity with the corresponding temporal filters and summed over all neurons:

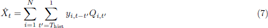

Because SAILnet only predicts deviations from a baseline, the mean and variance of the original stimulus were added back to the decoded patches and the patches were ordered into one sound waveform to create the decoded signal: 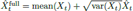.

### Quantification of Model Performance

To quantify our model’s performance, we calculated the pixel-wise mean squared difference between the original novel audio clip and the reconstructed audio clip. As described in the main text, to perform this comparison we first calculated the cross-correlation between the reconstructed signal and the original signal, and identified the time point at which the two signals had the highest correlation (representative example shown in Fig. 8A). We then aligned the two signals to be maximally correlated before computing the mean squared difference between them. This correction is necessary because the network does not naturally decode the phase offset of the original signal, and a naive mean squared difference calculation would predict that the untrained network has better performance than the trained networks due to destructive interference between the two signals (Fig. 9). The cross-correlation we calculate for this correction also served as a quantification for how similar the two signals are. In addition, we also quantified the mean squared difference between spectrograms of the original and reconstructed audios, as computed using Matlab’s spectogram function.

**Figure 8:**
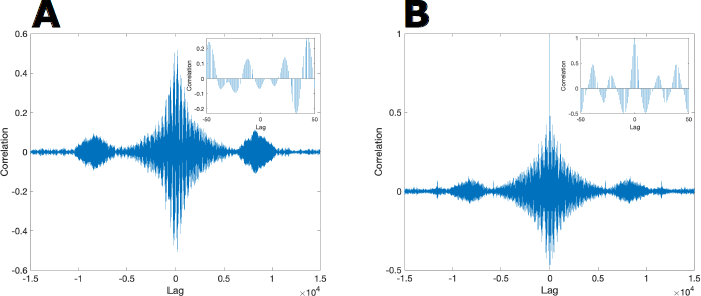
Correlations of novel signals. **A. Cross-correlation between a novel lo-fi audio clip and its reconstruction:** For each novel signal and the associated reconstructions, we compute the lag at which the correlation between the signals was the greatest; representative example using the output of a network that had been trained for 6,000 iterations is shown. There are notable oscillations in the correlation values at different lags, so we align the signals to have the greatest correlation before quantifying our network’s performance. **B. Auto-correlation of the novel signal.** The shape of this plot resembles the shape of the cross-correlation plot shown in (A), indicating similarity in structure of the novel audio and its reconstruction.

**Figure 9:**
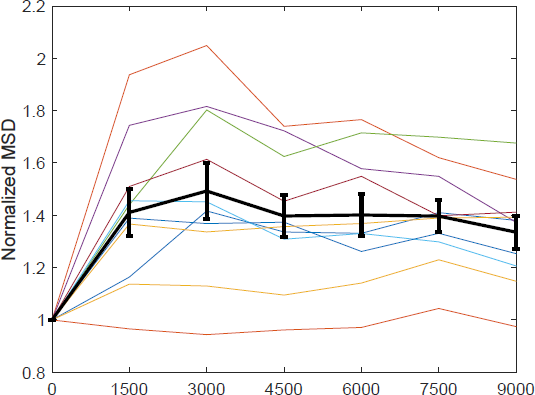
Point-wise mean squared difference between ten novel audio clips and their unshifted reconstructions. To quantify our model’s performance we calculate the point-wise mean squared difference between ten novel lo-fi audio clips and their raw reconstructions, without shifted the signals to be maximally correlated. Each colored line corresponds to a novel audio clip and the black line is the mean *±* SEM of the mean-squared error of the set. This metric erroneously indicates that the model did a better job of reconstructing the audio clip before any training has occurred, due to the fact that the reconstruction does not capture the absolute phase of a signal, and aligning of the signals is necessary.

**Figure 10:**
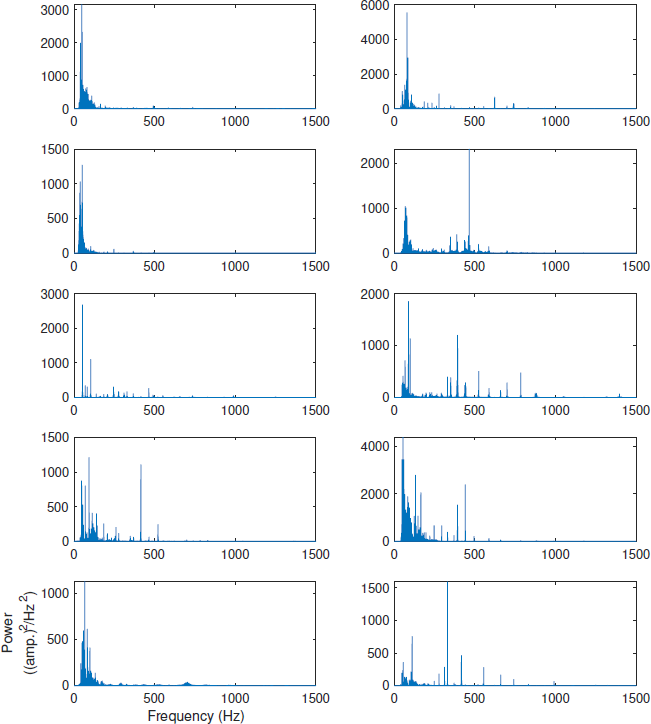
Power spectra of lo-fi songs. We obtained the ten lo-fi songs used to train our network from royalty-free music sites including pixabay.com, tunetank.com, chosic.com, and free-stock-music.com. The power spectra above show the frequency components of the lo-fi songs before any downsampling or normalizations were done. The power for all frequencies above 1,500 Hz was *∼*0. There are no significant frequency components above 1,000 Hz in any of the lo-fi songs.

**Figure 11:**
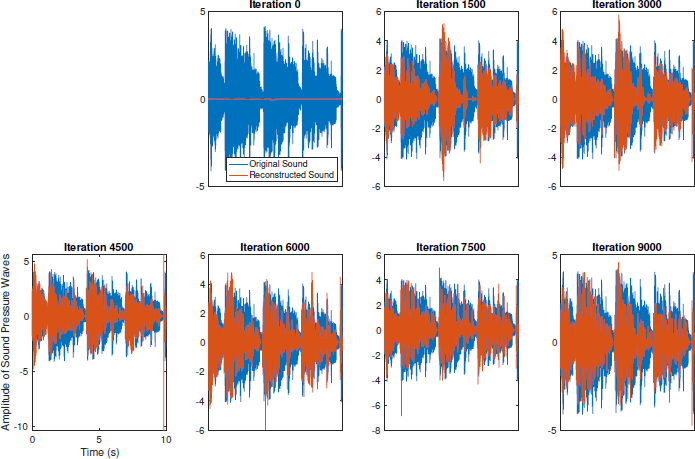
Sound waveforms of a novel lo-fi audio sound clip and the reconstruction of that clip. We use a lo-fi audio from pixabay.com as our novel audio for this example, which is not a part of the training set. The waveform of this audio after downsampling is shown in blue. Following training, this audio is reconstructed six times using the model parameters from every 1500 iterations. The waveform of the reconstructed audio is shown in red in the above examples. Despite the fact that lowest point-wise mean squared difference between the signals occurs before training if we do not shift the signals to be maximally correlated (Fig. 9), the waveforms of the reconstructed sounds more closely resemble the original waveform following training.

**Figure 12:**
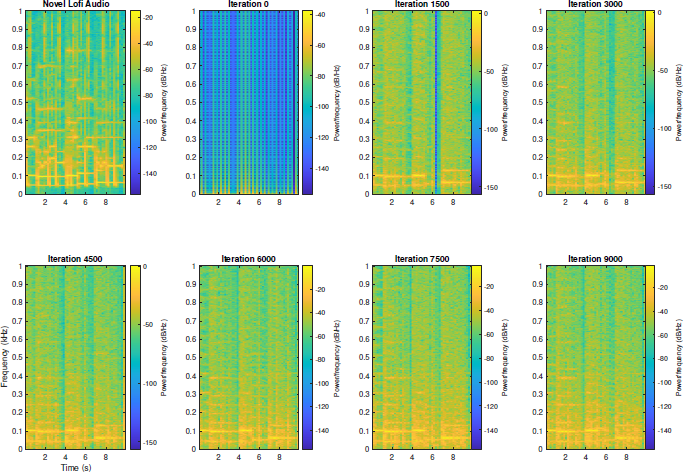
Spectrograms of a novel lo-fi audio and reconstructions of that audio. We use a lo-fi audio from pixabay.com as our novel audio for this example. Here we show the spectrogram of this novel audio (top, left), the spectrogram of the reconstruction before any training (top, second from left), and reconstructions of this sound after training. Reconstructions after training are shown for every 1,500 iterations of training. The low frequency components, up to 200 Hz, are captured best in all reconstructions done after at least 3,000 training iterations.

## Figures

Figures created in MATLAB were saved using the export_fig script, available on GitHub [26].

## Supplementary Files

1. Audio file of an exemplary neuron’s receptive field with a preferred frequency of 45 Hz.
2. Audio file of an exemplary neuron’s receptive field with a preferred frequency of 95 Hz.
3. Audio file of an exemplary neuron’s receptive field with a preferred frequency of 105 Hz.
4. Audio file of an exemplary neuron’s receptive field with a preferred frequency of 185 Hz.
5. Audio file of an exemplary neuron’s receptive field with a preferred frequency of 330 Hz.
6. **(a)** Novel lofi audio file taken from pixabay.com, a royalty-free music website. This song was originally 1 minute 55 seconds long and sampled at 48,000 Hz. The clip included in this file contains 10 seconds of that song downsampled to 2,000 Hz. **(b)** Reconstruction of the audio file in (a) using our network after 6,000 iterations of training.

